# Selection, Evolution and Persistence of Paleoecological Systems

**DOI:** 10.1101/2024.11.14.623653

**Authors:** Peter D. Roopnarine

## Abstract

The Phanerozoic fossil record can be organized as a nested set of persistent paleoecological units, ranging from paleocommunities to the Evolutionary Faunas. This paper argues that the basis for ecological persistence on geological timescales is rooted in the robustness of ecological communities, that is, the resistance and resilience of communities when perturbed by the environment. Here I present the Ecological Functional Networks Hypothesis (EFNH) that proposes that networks of species functions, or Ecological Functional Networks (EFNs), underlie ecological stasis and persistence, and that EFNs are both subject to selection and evolve. An EFN varies if the species composition and hence functional structures of its constituent communities vary, and EFNs may differ from each other based on the robustness of those constituent communities, numerical representation, and biogeographic distribution. That variation is subject to selection acting on EFN community composition, and determines both the persistence of an EFN and the differential persistence among multiple EFNs. Selection pressures on EFNs in turn exert top-down influence on species evolution and extinction. Evidence is presented to both establish the reality of EFNs in the fossil record, for example community structures that persist even as species composition changes, and the selection of EFNs, which is apparent during and after episodes of severe biotic turnover such as mass extinctions. Finally, tests are suggested that make the EFNH falsifiable, including testing the correlation between EFNs or EFN emergent traits and geological persistence, and using models of paleocommunity dynamics to examine the relationship between community or EFN robustness and geological persistence. The tests should be applied broadly throughout the Phanerozoic and diverse environments. The EFNH is part of a growing body of hypotheses that address multilevel selection and evolution of non-reproducing systems, including ecosystems and entire biospheres, and addresses those concepts on geological timescales.

## 3 Introduction

The Phanerozoic fossil record comprises hierarchically structured, multi-taxon, temporally bound, and compositionally persistent biotic units [Roopnarine and Banker, 2021]. Its diversity of multicellular lineages has persisted for ∼ 600 million years (my), standing in stark contrast to the microbially-dominated preceding ∼ 3 billion years. Persistent units encompass communities that vary spatially and temporally in species and phylogenetic composition, but that maintain a system of taxon functions, the functional networks formed from interactions of those functions, and the processes that emerge from the interactions. Persistence itself is a function of the robustness of a network or system to an ever-varying environment. In this essay I will outline a hypothesis, the Ecological Functional Network Hypothesis (EFNH), wherein networks of interacting ecological functions are the fundamental units from which paleobiotic persistence is derived. EFNH proposes that ecological persistence is selected and evolved, with selection acting on emergent properties of ecological functional networks (EFNs) that affect their ecological robustness. Species traits, biotic interactions, ecosystem biotic-abiotic interactions, and feedback within and among these, are ultimately responsible for the emergent properties that are subject to selection, but these properties, being emergent, are not reducible to individual species, nor functionally narrow phylogenetic lineages. The emergent properties change as community composition changes because of species evolution, extinction, and migration, but species evolution itself is subject to top-down feedback from community and ecosystem levels of organization and their dynamics [Roopnarine and Angielczyk, 2016]. Even as species are major contributors to system robustness, their own evolution is constrained or facilitated by EFN dynamics.

The EFNH is presented as follows. First, persistent ecological functional networks are defined, and evidence presented to establish their reality. Second, mechanisms and supporting evidence are proposed to explain how EFNs arise via multi-level selection, and tests are discussed that could falsify the hypothesis. Finally, the relationship of EFNH to other proposals of ecological system selection, evolution and persistence are discussed, as well as additional hypotheses of multilevel system selection, evolution and persistence.

## 4 Evidence

Scientific hypotheses are proposed to address empirical observations that lie outside the domain of, or conflict with, current theory. The observations of concern here are paleoecological phenomena, including: (1) a nested hierarchy of geologically persistent paleoecological assemblages; (2) paleoecological communities characterized by morphologically or taxonomically static species; and (3) paleoecological functional frameworks that are geologically persistent even as species composition within a framework changes.

### 4.1 Persistent systems

At the heart of the EFNH are networks of biotically interacting ecological functions, defined by species functional traits, such as body size, functional morphologies, and biogeochemical activities. It is now commonplace in paleoecology to characterize taxon assemblages and paleocommunities according to functional traits at both taxon and community levels–e.g. functional disparity– [Dineen et al., 2019, Cole and Hopkins, 2021] where the measures are often emergent properties. Paleocommunities are systems, with characteristics depending ultimately on the properties and relationships of constituent species, and functionally similar or redundant species, their interactions with the external environment, and the outcome of the feedback of system dynamics to constituent entities.

EFNs are one level in an ecological hierarchy that ranges from individual organisms to local populations or avatars, to local communities and metacommunities [Damuth, 1985, Eldredge, 1989]. The hierarchy occupies a dimension of spatial contemporaneity, encompassing population distributions and both species and community spatial connectivity, but there is also a temporal dimension that describes geological persistence. A significantly persistent paleoecological unit is one whose geological duration exceeds that of the transitional interval that separates it from preceding and succeeding intervals of the same type. Sepkoski’s Phanerozoic Evolutionary Faunas [Sepkoski, 1981] and Boucout and Sheehan’s Ecologic Evolutionary Units (EEUs) [Boucot, 1983, Sheehan, 1996], are canonical, inclusive, and rigorously defined units at the top of the hierarchy. The Evolutionary Faunas group Phanerozoic marine animal families and genera according to their ecological characteristics and origination-diversification-extinction histories [Alroy, 2004], and are separated by intervals of major turnover and extinction. Nested within the Sepkoskian Faunas are the EEUs, each comprising contemporaneous benthic marine communities of similar ecological structure, much of which is generated by persistent phylogenetic lineages. EEUs vary in duration from 30-140 my, separated by much shorter transitional intervals of 3-5 my, which are often associated with times of increased extinction. Species composition varies within EEUs, but compositional stability at higher taxonomic levels suggests significant ecological continuity, to the extent to which ecological functions are conserved within a particular lineage. This stratigraphically constrained variability of taxon composition is common in the Phanerozoic record [DiMichele et al., 2004], including Permian tetrapod terrestrial faunas [Olson, 1952], late Paleozoic terrestrial floras [DiMichele et al., 2002, Willard et al., 2007], and both late Cenozoic terrestrial mammalian assemblages [Barry et al., 2002] and tropical coral reefs [Pandolfi and Jackson, 2006]. An EFN is more constrained because continuity of both functions and the network of interactions are required. Functions arise and disappear according to the persistence of the species performing them, but accordingly can recur discontinuously and be performed by phylogenetically distant species [Banker et al., 2022]. An EFN is therefore the system of functions and their pattern of interaction, regardless of species composition.

#### 4.1.1 Paleocommunity persistence

Despite a “haziness” inherent in the delineation of ecological communities [Yodzis, 1988], species biotic inter-dependencies, and environmental requirements do give rise to replicable species assemblages and networks of interactions. A classical ecology argument arises from contending views of communities as highly integrated and somewhat functionally inflexible species assemblages [Clements, 1916], versus more random assemblages of species based largely on environmental requirements [Gleason, 1926]. Yet replicability of species assemblages and networks of interaction, when extended to the fossil record and the temporal dimension, are the main bases for paleocommunity recognition.

Brett and others [Morris et al., 1995, Brett et al., 1996] documented species-level compositional stability in mid-Paleozoic faunas from the U.S. Appalachian Basin, with units persistent for 3-7 my. The proposed Coordinated Stasis hypothesis postulates that the persistence of morphologically static species assemblages, and their synchronous turnovers, are driven by either rigid patterns of biotic dependencies (“ecological locking”), or environmental tracking. Ecological locking implies strong interspecific interactions, that is, individuals have significant per capita effects on the populations of other species. Theoretical ecology, however, predicts strong interactions to have destabilizing effects on community structure [May, 1972, McCann et al., 1998], and one would least expect ecological stasis under such conditions. The Appalachian Basin assemblages do exhibit very little change, and may be a limiting case of a more general phenomenon of ecological stasis, wherein taxon composition varies even as emergent community features remain stable. Ecological communities are rarely spatio-temporally homogeneous, but instead occupy variable environments that influence both species composition and therefore community functional structures. Compositional variation is therefore not surprising, but functional persistence is expected when that variation is phylogenetically constrained and within-lineage ecology is conserved. For example, Silurian brachiopod assemblages exhibited changes of species composition over ∼ 30 my, but species richness and ecological characteristics changed little [Watkins et al., 2000]. Functional persistence is expected even when compositional variation is phylogenetically broad but multiple lineages in an assemblage share environmental requirements. For example, bivalve assemblages of the Jurassic U.S. Western Interior exhibited moderate compositional stability, but with significant amounts of turnover among stratigraphically successive units with durations ranging from 2-6 my [Tang and Bottjer, 1996]. Thus there is an organizational hierarchy of community temporal compositional stability. This stability may extend beyond species composition to other community characteristics, e.g. bird richness on the island of Hanimaluoto off Finland remained nearly constant between censuses taken over 50 years (1967-2013), despite significant turnover of species composition [Hanski, 2019]. Modern beetle assemblages from southern England compared to those dating to 43,000 years ago, and therefore separated by a glacial interval, exhibited moderate amounts of species turnover, but constancy of species richness [Hanski, 2019].

### 4.2 Ecological functional networks

An EFN is a system consisting of ecological functions, performed by species, that interact with each other. “Function” is used broadly here to refer to any action that members of a species undertake to ensure individual or reproductive success, and that has an impact on other individuals, including those belonging to other species. Examples include foraging, ecosystem engineering, and actions affecting biogeochemical cycles, such as microbial decomposition. Functions affect other functions, either facilitating or inhibiting them. For example, giant kelp that attain great height and form a canopy at the water’s surface to perform the vital role of photosynthesis and production, also dampen hydrodynamic forces thereby promoting local biodiversity, and shade the benthic understory thus inhibiting benthic primary productivity [Detmer et al., 2021]. Multiple species may perform a function redundantly, but those species are unlikely to be completely redundant because they differ in other traits, functions and environmental requirements. This is a key mechanism of how biodiversity promotes community robustness. Therefore a function, as used in the current context, can be an abstraction of one or more species, and an EFN is not a system of species, but a system of abstracted functions. Recent work on Miocene-Pleistocene terrestrial communities of the Iberian peninsula showed that their functional compositions, and networks of functional interactions, persisted in the face of species opportunistic and climate-influenced immigration and extinction [Blanco et al., 2021]. Network modules of species assemblages persisted an average of 0.9 my, modules defined on the basis of functions persisted ∼ 2.8 my, and three networks of functional modules persisted 2.58, 4.66 and 9.37 my. Functional structures were therefore more persistent than species, and species immigrating into an EFN generally performed roles already present in the EFN. Networks of functional modules typify EFNs, another example being late Permian networks of southern African terrestrial communities, which although spatio-temporally variable in taxon composition, were structurally cohesive for more than 10 my [Roopnarine et al., 2017], changing only during the Permian-Triassic mass extinction (PTME). Early Triassic successor communities represented new and structurally distinct EFNs.

Therefore, phylogenetic change and species turnover occur in EFNs, but community structure and processes are continuous. Taxon stability in an EFN is unsurprising if new species are descendants of earlier species in the EFN and retain ancestral functional traits, but taxonomic variability among communities can be great enough to generalize an EFN as a set of structurally similar, but compositionally varying communities. Modern tropical coral reefs are an example, with a diverse set of scleractinian taxa forming physiologically and ecologically similar systems, while varying in terms of species richness, composition and dominant species. This is a somewhat Clementsian view [Clements, 1916] of communities, but at a level of organization above that of individual species, with the focal units being species functions that interact with other species functions. Thus, it is the integrated systems of functions that persist, and not necessarily systems of particular species.

The types of organisms, environmental conditions, and ecosystems vary across this range of examples, but a common theme of geological persistence holds. In some cases, persistence is apparently limited to a subset of what must have been a series of communities, e.g. floras, and in other cases there is broader ecological continuity. Timescales also span a range of durations, from tens of millenia to millions of years. Yet what is noted is the apparent persistence of interacting ecological functions, and by extension community dynamics, because of the presumed conservation of ecologies among diverse sets of taxa.

## 5 Selection and evolution

EFNH proposes that: (1) EFNs belong to a class of entities above the species level, that includes clades, ecosystems and function interaction networks [Doolittle and Inkpen, 2018, Papale and Doolittle, 2024]; (2) that geological persistence arises from selection acting on EFN properties which vary according to variance within and among communities; and (3) that EFNs represent evolved functional networks. According to Hull’s replicator-interactor framework of evolution by natural selection [Hull, 1980], interaction with the environment may take place at a level of organization above that of replicators, with replicators causing the interaction which in turn affects replication. EFNs are interactors, and species are both interactors and replicators. It is unnecessary, however, for EFNs to be replicating units with heritable traits to evolve, for selection here is not causing differential reproductive success, but instead differential persistence [Dussault and Bouchard, 2017]. It is also unnecessary for EFN-environment interactions to be reducible to the species level, for there are emergent EFN properties that are subject to selection by the environment, the outcome of which will feedback to species replication and evolution. EFNH further proposes that feedback constrains both functional innovation and divergence of species within an EFN, as well as the success of species introduced into an EFN, thus extending EFN persistence even if species composition varies. There are subsequently several challenges that EFNH must address, including: (1) identifying EFN traits acted upon by selection; (2) explaining how the concept differs from lineage-based evolution by natural selection; and (3) outlining how EFNH explains geological persistence.

### 5.1 Emergent traits

An emergent system trait is one that is neither shared with, nor reducible to the traits of any single entity within the system. EFNH requires that EFNs be characterized by emergent traits and not the functional traits of individual species in any particular community of an EFN, for in that case any proposed stasis, persistence or selection of an EFN would be indistinguishable from those processes acting at the species level. EFN traits must therefore be emergent, above the species level. The traits are rooted ultimately in the emergent properties of an EFN’s constituent communities, in the same manner in which a community’s emergent traits are rooted in the traits of its constituent species, but similarly, an EFN will also exhibit traits that are not reducible to the level of any of its constituent communities, unless it consists of a single community.

Emergent community properties are one class of EFN traits, and include properties of individual communities such as total abundance, geographic distribution, ecosystem functions and community robustness (resistance, resilience and anti-fragility [Taleb, 2014, Munoz et al., 2022]). Both ecosystem functions and robustness are dependent on the emergent functional networks formed by species. It is important to distinguish those community emergent properties that are variable among communities that share an EFN, from properties that emerge collectively from all the communities sharing an EFN, and that therefore may distinguish among communities. Examples of the former are exhibited by modern coral reefs. Modern reefs are all descended from communities that arose from the earliest scleractinians in the Middle Triassic, and they share later important events, such as a late Neogene parrotfish (family Labridae) diversification in coral reefs [Choat et al., 2012], as well as both environmental requirements and physiographic impacts. In broad functional structure, modern reefs may all be assigned to a single EFN on the basis of shared history and functional structure, but are also distinguishable on the basis of other characteristics that are important to persistence. These include species richness, geographic distribution, and resilience founded on functional diversity. Among tropical western Atlantic reefs one can recognize differences of richness and composition between the Bahamas, the Gulf of Mexico, Greater Antilles, Lesser Antilles, and South America. There are also geologically persistent variants and variants of shorter duration [O’Dea et al., 2020]. A second class of EFN traits are cumulative emergent properties that are not community properties, but rather result from sets of communities, including the number of communities that share an EFN (e.g. the number of tropical western Atlantic coral reefs) and their geographic distribution.

Emergent EFN properties also serve to distinguish among EFNs. For example, forests and grasslands, which often occupy the same geographic landscape as alternative ecological terrestrial ecosystem states, have at any given time and under specific environmental conditions different emergent properties. Those properties include the number of communities (forest versus grassland), their richnesses, robustness against perturbations such as drought and fire, and different agents, rates and mass transfers within biogeochemical cycles. Particular times and places may favour one EFN over another.

Thus there is a nested hierarchy of emergent properties that characterize the hierarchy of ecological units. Species have properties that are not reducible to individuals or populations, communities have properties that are not reducible to their member species but nonetheless can affect community persistence, and EFNs have properties that are not reducible to the community level and which are important to the persistence of the EFN. The interactions between these properties and environmental perturbations form the basis for selection leading to differential persistence, and selection at one level may affect emergent properties at a higher level. This is a central concept in Lenton et al.’s [Lenton et al., 2021] “survival of the systems”, and Dussault and Bouchard’s Persistence Enhancing Propensity [Dussault and Bouchard, 2017]. Selection acting on a population affects the geographic distribution of the species, which in turn can affect community robustness depending on the presence or absence of the species or its abundance, and subsequently affects community persistence and hence the number and variability of communities within an EFN. The interactions of emergent properties at a higher level in turn feeds back to entities at lower levels because: (1) the number of communities within an EFN, an emergent property, can affect community persistence if individuals disperse among communities; and (2) community robustness can affect the survivability of member species. Ultimately the success of species and communities depends on selection acting on species traits and functions, but both community and EFN emergent properties are critical to species persistence and longevity.

### 5.2 EFN selection and evolution

There are several classes of evolutionary processes that occur at multiple levels within an EFN. First, species functions evolve according to the conventional framework of selection acting upon heritable traits and their variation, giving rise to genetic lineages [Lewontin, 1970]. Second, species persistence and hence evolutionary success, may depend on the completion and persistence of emergent community processes [Dussault and Bouchard, 2017, Lenton et al., 2021], and the system’s capacity to withstand (resistance), recover from (resilience), or thrive (anti-fragility) when perturbed. Third, community persistence is dependent upon both the success of its constituent entities, and the success of its emergent processes and robustness. Fourth, the success and persistence of an EFN depends both upon its numerical representation, that is the number of communities sharing the EFN, which must exceed zero, and the persistences of those communities. But it isn’t necessary for the number of communities to exceed one for an EFN to be persistent.

Selection acting on EFN properties explains geological persistence of higher-level paleoecological units. Here I use the familiar heuristic of a system state landscape to explain EFN selection (Figure 1). The landscape’s plane depicts a multidimensional space of EFN emergent properties occupied by communities. EFN structure, functional groups, network topology and interaction parameters, determine community position on the landscape. Elevation represents the likelihood of EFN and therefore community persistence, ℒ (*P*), and peaks on the landscape are therefore local maxima. Multiple peaks represent multiple EFNs, with distance between peaks, a landscape metric, measuring differences of functional composition and network topology (Figure 1a). There can be multiple communities at any given time clustered around a peak, each representing a community consistent with the EFN, but potentially differing from others in the cluster because of species-level variation such as abundance, assemblage composition, and community-level properties such as robustness. Under perturbative conditions, these variants of the EFN are expected to exhibit variable persistence.

**Figure 1:**
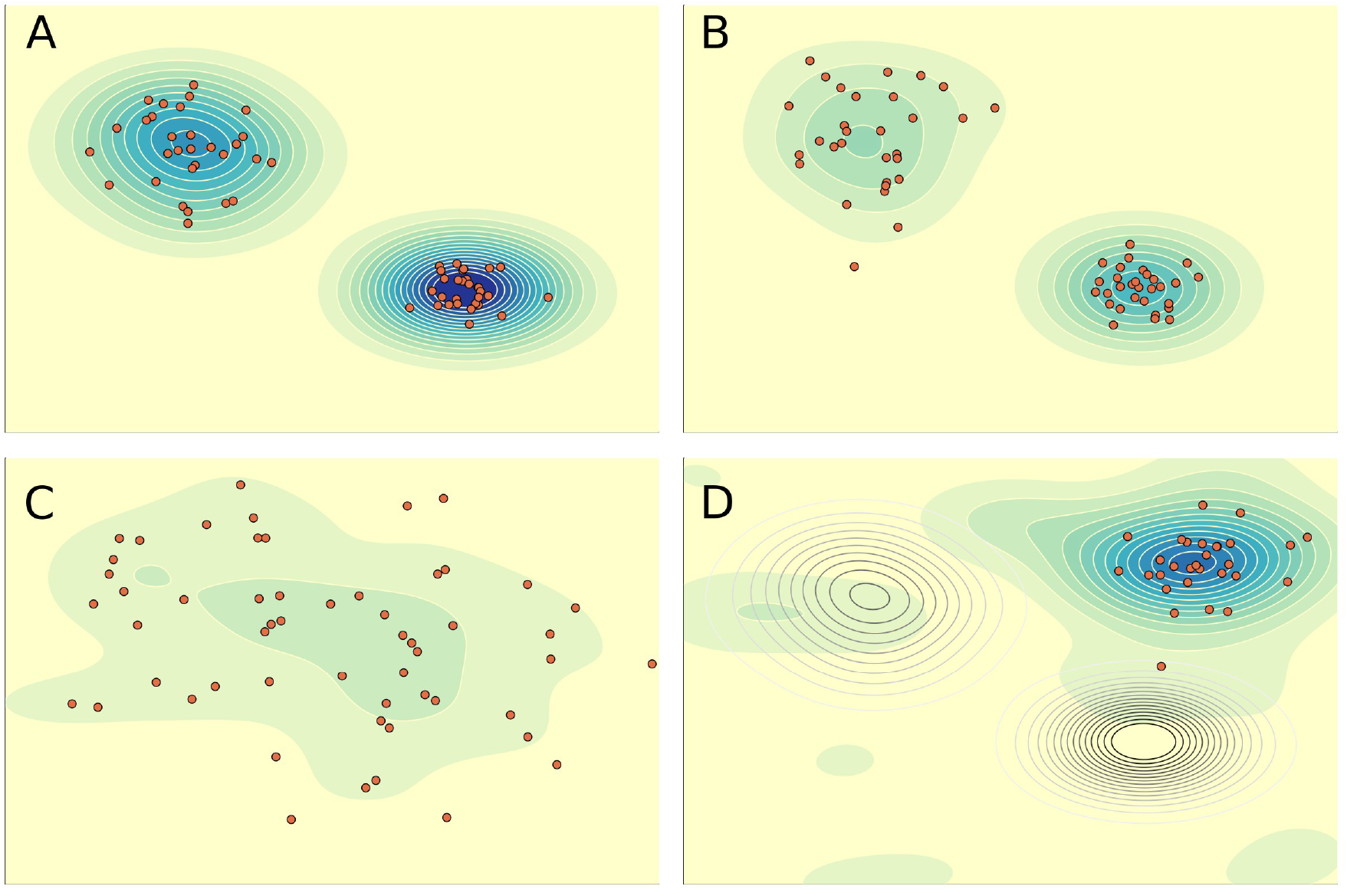
EFN heuristic state landscape. Contours measure the likelihood of persistence, ℒ (*P*) (bluer colors = greater ℒ (*P*)). Symbols (red) are individual communities. **(A)** There are two distinct EFNs, each represented by a cluster of communities. **(B)** Perturbed conditions reduce ℒ (*P*) of both EFNs. **(C)** EFNs are destroyed by high magnitude perturbations, resulting in a high variance of communities, and landscape exploration. **(D)** A new EFN eventually arises. Unfilled contours show locations of previous EFNs (**A**).

Landscape topology is dynamic, changing and varying over time, driven by the frequency and magnitude of perturbation regimes. The landscape is not a zero sum space, as there is no empirical evidence supporting zero sum constraints directing the history of life. Similarly, EFN persistence and the co-existence of multiple EFNs do not reflect steady state conditions, for these are thermodynamically open systems operating far from equilibrium, and constantly responding biogeographically, ecologically and evolutionarily to a dynamic environment. Reduced perturbation or frequency increases the ℒ (*P*) of multiple community variants, and lesses the differential gradient between EFNs, potentially allowing a greater number of EFNs to co-exist, and increasing global functional diversity (Figure 1a). Such conditions also mark times of reduced top-down constraints of EFN dynamics on species evolution, and the occurrence of geologically persistent EFNs. Increasingly severe perturbative conditions degrade gradients around landscape peaks, reducing ℒ (*P*) of variants around the peaks (Figure 1b). These are times of ecological crisis marked by species extinctions, and the collapse of communities and ecosystems. The energetic power of a perturbation, if great enough, will reset the landscape, eliminating EFNs on geologically or even ecologically short timescales, e.g. the Chicxulub impact at the end of the Cretaceous.

#### 5.2.1 Evidence of EFN selection

Less powerful or extended perturbations, which can nonetheless have severe cumulative effects, e.g. the Permian-Triassic mass extinction (PTME), offer opportunities to examine landscape dynamics and EFN selection. For example, taxonomic extinction and EFN collapse were decoupled during the PTME [Roopnarine et al., 2019, Huang et al., 2023]. The transition was marked by species extinctions that either occurred in multiple phases, or extended through the PTME, whereas EFN collapse, marked by the collapse of guild-level trophic networks, occurred toward the end of the PTME. Functional redundancy and the semi-independence of EFN robustness and species composition resulted in EFN persistence beyond the extinction of multiple constituent species. The Late Ordovician mass extinction (LOME) displays a similar pattern [Droser et al., 2000]. EFN selection occurs under these circumstances. Consider a single EFN. Community variants have different ℒ (*P*), either because species or functions held in common vary in details, or because they differ compositionally while belonging to the same EFN type. Compositional variation and differential species vulnerabilities may lead to extinction of some EFN variants and the survival of others. Multiple EFNs subjected to the same dynamic circumstances would respond differently if they differ in relevant emergent properties. Some EFNs therefore could retain greater numbers of variants than others, thus increasing their numerical representation, or an EFN might disappear altogether. Both scenarios are consistent with proposed systems-level selection mechanisms, including increased numerical representation, and persistence-based selection [Doolittle, 2014, Dussault and Bouchard, 2017].

Short-lived paleoecological units are observed to separate those of longer duration, an observation consistent with EFNH, which predicts that boundaries betwwen EEUs, marked by significant taxonomic and paleocological changes, also mark changing EFN landscapes. EFN destruction during environmentally challenging times releases species from top-down constraints on evolution and functional structure. What follows is an interval of landscape exploration by surviving and perhaps re-diversifying lineages, and communities that have no antecedents in the previous EFN (Figure 1c), a process that has been referred to as “random rewiring” [Lenton et al., 2021]. These aftermath communities are predicted to be short-lived, not because of ecological instability, but because there are numerous alternative ways in which the evolving lineages can assemble into more efficient and robust EFNs. They are likely to do so incrementally via species coevolution [Roopnarine and Angielczyk, 2016], facilitating some continuity of system information [Lenton et al., 2021]. Probable examples include LOME and PTME aftermaths. Counterfactual models of robustness of Early Triassic EFNs from the Karoo Basin of South Africa showed precisely this, with counterfactual EFNs being significantly more robust than the actual EFN [Roopnarine and Angielczyk, 2015, Roopnarine et al., 2019]. The subsequent return of long persistent EFNs indicates selection of emergent properties, leading to increased ℒ (*P*) and the resumption of top-down feedback to species selection.

Another potential line of evidence crops up in community food web networks. One property of those networks, and all other networks with directional links between network nodes, is the node in-degree distribution, Pr(*k*). Such links point away from resources or prey species, and toward consumers, indicating pathways of energy transfer through the community. Measures of Pr(*k*) for numerous modern food webs ranging in size from dozens to hundreds of species [Dunne et al., 2002, Roopnarine and Hertog, 2013], and encompassing a variety of terrestrial and aquatic habitats, all share a notable feature; the proportion of species of low in-degree, or trophic specialists, declines as in-degree increases, leading to communities comprising greater numbers of relatively specialized consumers versus a lesser number of relatively generalist consumers. Pr(*k*) is frequently consistent with exponential or power law distributions, referred to here collectively as “decay distributions”, with the rate of decline varying from exponential to slower than exponential. Having a decay distribution where more species tend toward trophic specialization than generalism, suggests the following underlying eco-evolutionary mechanisms: (1) Species might tend toward specialization because of selection for a more efficient utilization of resources, or for minimizing interspecific competition; (2) trophic generalists might be rarer because efficient generalism requires a broader array of functional traits; or (3) generalist predators require greater power in order to utilize a more functionally diverse set of resources [Vermeij, 2023]. Logically, stenotopic specialists would have higher probabilities of extirpation or extinction if trophic chains are disrupted because they have fewer resources, thereby suggesting that the evolution of a decay-type in-degree distribution would require an evolutionarily significant interval of relative calm on the EFN landscape.

An alternative explanation that is consistent with EFN selection and evolution comes from modern network theory. A key finding there is that the link distributions of real networks, e.g. the Internet, transportation networks and food webs, deviate strongly from random networks (e.g. Poisson or normal distributions) and instead tend to be decay distributions. Networks with such distributions are topologically robust against the random removal (extinction) of nodes or links. For example, because most Internet servers are linked to only a few other computers (specialists), the loss of a single server will on average have no noticeable impact on the functioning of the Internet. The same would hold true for the random removal of a species from a sparsely connected food web in which species are predominantly specialized [Roopnarine, 2006, Dunne et al., 2002]. In contrast, the loss of a highly connected server or species could have system-wide effects. The connection to EFNH arises when we consider how different types of networks come to have decay degree distributions. Whereas the robustness of artificial networks often arises from intentional design [e.g., Mozafari and Khansari, 2019], preferential attachment (“the rich get richer” [D’souza et al., 2007]), or iterative correction of link distributions in response to negative perturbations [e.g., Yang et al., 2015], ecological community robustness is less easily explained. The distributions in modern communities could have arisen either fortuitously as a result of species-level eco-evolutionary constraints and opportunities as described above, in which case community robustness would not be a factor as specialist species would be more prone to extinction, or by the iterative evolution of the communities and distributions themselves. The latter would imply: (1) a differentiation of ℒ (*P*) between communities on the basis of disturbance histories and link distributions, and (2) the existence of top-down feedback from system dynamics to species-level evolution [Roopnarine and Angielczyk, 2016, 2012], because too many generalists in a community would increase its vulnerability to propagating perturbations, and the potential for both species extinction and system collapse. A differentiating factor among communities would therefore be both robustness against perturbations to species, and the subsequent persistence of a network structure on evolutionary and geological timescales.

### 5.3 Testing the EFNH

EFNH is consistent with a range of observed phenomena, but must be falsifiable. Two tests are suggested: (1) constructing community functional networks throughout the Phanerozoic; and (2) testing predictions with models of paleocommunity dynamics. The tests require construction of networks of taxa aggregated according to function, with links corresponding to classes of biotic interactions, and interactions between groups and the environment. EFNH predicts a positive correlation between system persistence and the presence of decay in-degree food web distributions. EFNs also may be suitably represented by hypergraphs, where species are nodes or vertices as in conventional network representations, but where (hyper)edges represent multidimensional relationships and interactions among species. Given the divisibility of Earth’s history into intervals characterizable by particular geophysical and geochemical conditions, secular trends or transitions such as continental configurations and climatic conditions, and biological increases of organizational complexity, body size, metabolic rates and power, colonization of ecospaces, and both ecosystem and Earth system engineering, EFNH predicts that at any given time multiple communities will share hypergraph properties, such as motifs [e.g., Contisciani et al., 2022, Lotito et al., 2022], and potentially be isomorphic when species are abstracted as functions [Feng et al., 2024]. Tests of this nature will rely heavily on the continuing assembling of paleocommmunity data and functional interpretation. EFNH further predicts that paleocommunities that share the characteristics of an EFN will do so during the same interval of time regardless of taxon variation. Such uniformity will disappear during transitional intervals, perhaps being replaced by communities of more singular character. Furthermore, if transitory intervals have sufficient stratigraphic resolution, taxon turnover and extinction will precede final EFN breakdown, as already described in earlier examples.

The second test examines EFN robustness using models of community dynamics and perturbation. Current network models of cascading secondary extinctions chronically underestimate dynamic impacts because they lack feedback processes [Dunne and Williams, 2009, Bodini et al., 2009], whereas available feedback models can be improved with incorporation of expanded species functional traits [Roopnarine et al., 2017, Huang et al., 2023]. Suitable models that simulate community dynamics require parameters that are difficult to gather or infer for fossil taxa, such as population sizes [but see Marshall et al., 2021], intrinsic rates of population increase, mortality rates, and interaction strengths, but recent advances that rely largely on scaling relationships of modern taxa [Brown et al., 2004, Savage et al., 2004] and uniformitarian assumptions, plus increasingly sophisticated inferences from fossils themselves, place enhanced models well within reach.

Ultimately, EFN structure, whether represented as ordinations of functional traits, conventional networks, or hypergraphs, may be combined with dynamic models to understand both the changing ℒ (*P*) of individual communities and an EFN as demonstrated by the following example. Roopnarine et al. [Roopnarine et al., 2022] presented a process-based model reconstructing the dynamics of giant kelp (*Macrocystis pyrifera*) forests of the North Pacific prior to the extinction of the megaherbivore Steller’s sea cow (*Hydrodamalis gigas*), with additional species including an understory alga (*Chondrocanthus corymbiferus*), the purple sea urchin (*Strongylocentrotus purpuratus*), the sea otter (*Enhyda lutris*) and the sunflower sea star (*Pycnopodia helianthoides*). This historical model was tested under conditions of sea star wasting disease, and the disease coupled with a persistent multi-annual ocean heat wave. Figure 2 presents the system EFN as hypergraphs of the model system under both perturbative conditions. Taxa in the model are united by three hyperedges that capture both predatory and competitive interactions. All the hyperedges are present under both perturbative scenarios, but the relative strengths of the interactions are reversed. This was interpreted as a decrease of robustness, and hence ℒ (*P*), under the combined perturbations of disease and warming (Figure 2b), both because of the removal of the sea star which is an important urchin predator, and the depression of kelp growth rates by increased water temperature.

**Figure 2:**
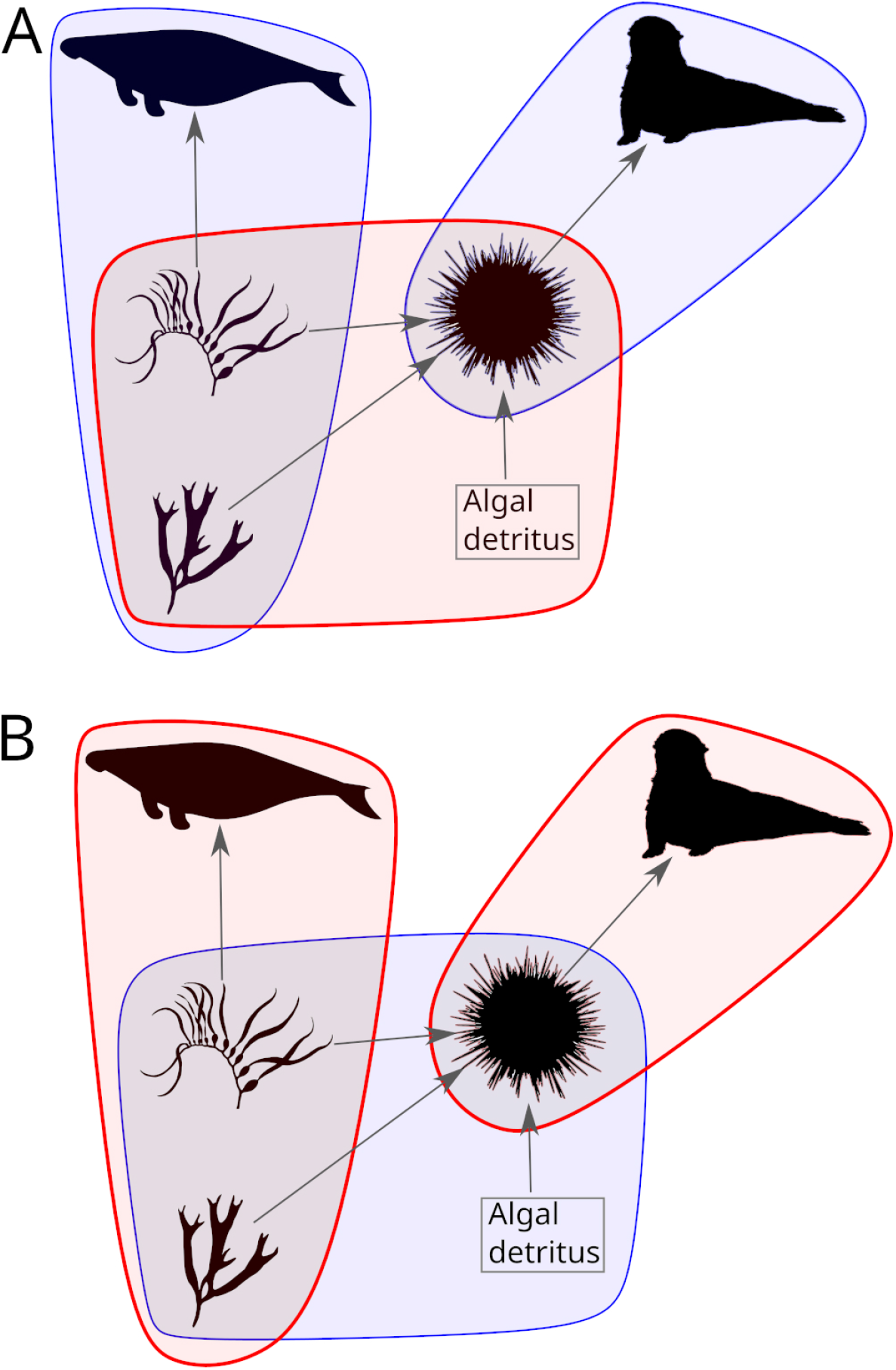
EFN hypergraphs of species functions and interactions in a model of historical North Pacific giant kelp forest ecosystems including the extinct megaherbivore *Hydrodamalis gigas*. Hypergraph edges connect two or more taxa (including algal detritus), with thick-outlined and red edges representing strong functional interactions in a particular community state, and thin-outlined blue edges representing weaker interactions. **A** Model EFN under conditions of sea star wasting disease, which has removed the urchin predator sunflower sea star *Pycnopodia gigas*. **B** EFN under conditions of both disease and a persistent marine heat wave. Note the reversal of edge strengths between the two EFN states. Gray arrows in both graphs, showing food web interactions, do not capture the full functionality of the system, ignoring competitive interactions and indirect relationships, e.g. via engineering impacts.

Finally, positive results of the above types of tests could be applied to conjectures that rates of background extinction have been steadily declining during the Phanerozoic. It has been suggested that the decline could result from changes to the ways in which perturbations propagate through communities because of Phanerozoic changes of community structures [Foote, 2000, Roopnarine, 2006]. Again, this would be a system-level phenomenon, offering a complimentary although not exclusive explanation from species-based evolution. The EFNH provides a legitimate mechanism.

## 6 Multi-level selection

In closing, EFNH is one of a growing number of hypotheses of multi-level selection. Those hypotheses argue for selection above the level of species, and address entities and processes in which species are constituent replicators, but wherein emergent properties of the higher levels are the interactors. Selection at higher levels feed back to lower levels, including species, influencing their evolutionary trajectories, and leading to differential system survival and greater persistence [Bouchard, 2008]. The nested hierarchy extends upward both materially and conceptually, because the organization, selection and evolution of systems are extendable to entire planetary systems, as in the Gaia hypothesis [Doolittle, 2014, Lenton et al., 2018], and ultimately to fine-tuned fundamental parameters of the Standard Model, as in the hypothesis of Cosmological Natural Selection [Smolin, 2004]. Emergent variation, system selection, and evolution likely represent a universal principle and historical mechanism that not only explain temporal change and stability on multiple timescales but, through multi-level feedbacks, unite multiple levels of material organization and complexity.

